# LungCancer3D: A Comprehensive Database for Integrating Lung Cancer Chromatin Architecture with Other Multi-omics

**DOI:** 10.1101/2021.10.26.466013

**Authors:** Xinyu Wu, Anlan Jiang, Jixin Wang, Shiyang Song, Yaping Xu, Qian Tang, Shirong Zhang, Bing Xia, Xueqin Chen, Shenglin Ma, Jian Liu

**Affiliations:** Zhejiang University-University of Edinburgh Institute (ZJU-UoE Institute), Zhejiang University School of Medicine, International Campus, Zhejiang University, Haining, 314400, China; Hangzhou Cancer Institution, Affiliated Hangzhou Cancer Hospital, Zhejiang University School of Medicine, Zhejiang University, Hangzhou, 310002, China; Affiliated Hangzhou First People’s Hospital, Zhejiang University School of Medicine, Zhejiang University, Hangzhou 310006, China

## Abstract

With the breakthrough of chromatin conformation capture technologies in recent years, the importance of three-dimensional (3D) genome structure in gene expression, cell function regulation, disease occurrence, and development has been gradually recognized. To provide a comprehensive visualization of chromatin architecture and other multi-omics data for lung cancer research, we have constructed a comprehensive database, LungCancer3D (http://www.lungcancer3d.net). This web-based tool focuses on displaying human lung cancer-related HiC data along with a variety of other publicly available data, such as RNA-seq, scRNA-seq, ATAC-seq, ChIP-seq, DNA methylation, DNA mutation, and copy number variations. Researchers can visualize these diverse multi-omics data directly through the genome browser and discover how the genes expression is regulated at diverse levels. For example, we have demonstrated that the high expression level of C-MYC in lung cancer may be caused by the distant enhancer introduced by the *de novo* chromatin loops in lung cancer cells to bind the C-MYC promoter. The integrated multi-omics analyses through the LungCancer3D website can reveal the mechanisms underlying lung cancer development and provide potential targets for lung cancer therapy.

## INTRODUCTION

Lung cancer continues to be the leading cause of cancer death worldwide [1]. Global Cancer Statistics 2020 showed an 11.4% incidence rate with a death rate of 18% worldwide [1]. Although targeted therapies for lung cancer, such as EGFR, have achieved good efficacy in a subset of patients, most patients still do not have targeted drugs for treatment [2]. Therefore, investigation of the mechanism of lung cancer development is urgently needed.

Various genomic and epigenomics studies have been utilized to reveal the mechanism in lung cancer progression, but the integration of multiple omics data is not effectively conducted [3]. Thus, many studies heavily depend on the limited omics data to understand lung cancer development. For example, it is still relatively difficult to comprehensively integrate multiple omics to study the development of lung cancer, especially the integration of the newly emerging HiC omics that detect chromatin interactions and reveal three-dimensional chromatin structure.

The abnormal 3D structure of chromatin has been shown to promote the progression of diseases, including cancer [4]. Previous research also showed that the chromatin loops in lung cancer tissues or lung cancer cells differed from normal lung tissue [5]. The formations of chromatin loops were reported to regulate gene expression [6]. And the chromatin accessibility and DNA methylation affect the formation of chromatin loops [7]. However, how loops regulated adjacent genes at the genome-wide level is mainly unclear, and how dysregulated chromatin loops contributed to cancer development remains elusive. Studying the 3D chromatin architecture may lead to a new point in cancer studies. For example, 3D chromatin architecture can be potential therapeutic targets by regulating the formation of loops regulating the critical gene expression [8]. Practical tools that display 3D chromatin architecture information with the integration of other omics are needed. Several online databases have been developed to visualize chromatin interactions, such as 3DIV and 3D Genome Browser [9, 10]. However, none of them can show the integrated analysis of multiple omics, especially on lung cancer, such as integrating the visualization HiC with other omics data and searching a gene of interest. To overcome current limitations, we developed a website database providing users with visualized multi-omics data, including gene expression, copy number variation, somatic mutation, miRNA, DNA methylation, chromatin open accessibility, and histone modification information (Figure 1). In addition, we also integrated single-cell RNA-seq (scRNA-seq) data for normal human lungs, lung cancer, and human lungs without or with COVID-19 infection. The differential analysis results between cancer (mainly lung cancer) and normal samples were also displayed. Using this comprehensive website database allows users to better utilize genomic and epigenomic data to generate a functional interpretation of the identified chromatin interactions or search for potential therapeutic targets for future research.

**Figure 1.**
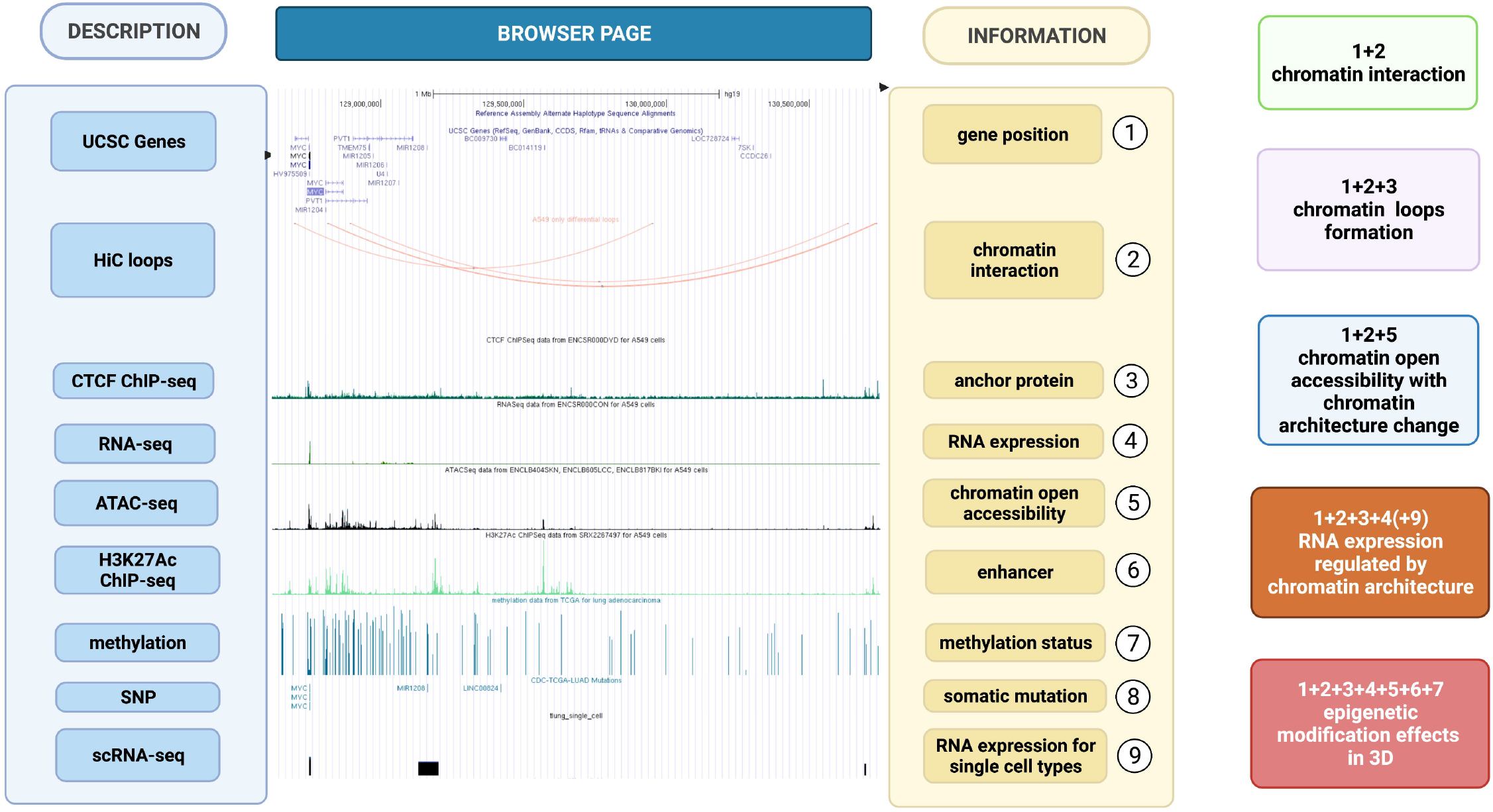
Overview of multi-omics data available at the website and examples of data integration. The left column shows the description of each track. The middle is the browser’s screenshot at the chromatin around lung-cancer-associated differential loops near the oncogene C-*MYC*. The right column is the functional interpretation of what each track can present. Users can utilize several combinations from different tracks to generate information related to chromatin interaction. (Created with BioRender.com)

## MATERIALS AND METHODS

### Website development and data source

Our registered website is www.lungcancer3d.net. UCSC mirror package was downloaded and installed on our web server by method browserSetup (genome browser in the cloud introduction) [11]. MariaDB was used on the backend to support the data structure. Apache HTTP server was used for cross-platform web services. BigWig file format is the basic file format recognized by MariaDB. Currently, our webpage only supports human genome assembly hg19, which was also downloaded from UCSC website. The raw data related to human lung cancer studies were downloaded from public databases and described in Table 1 [12] [13] [14] [15] [16] [17] [18] [19] [20] [21] [22] [23] [24] [25] [26] [27].

**Table 1.**
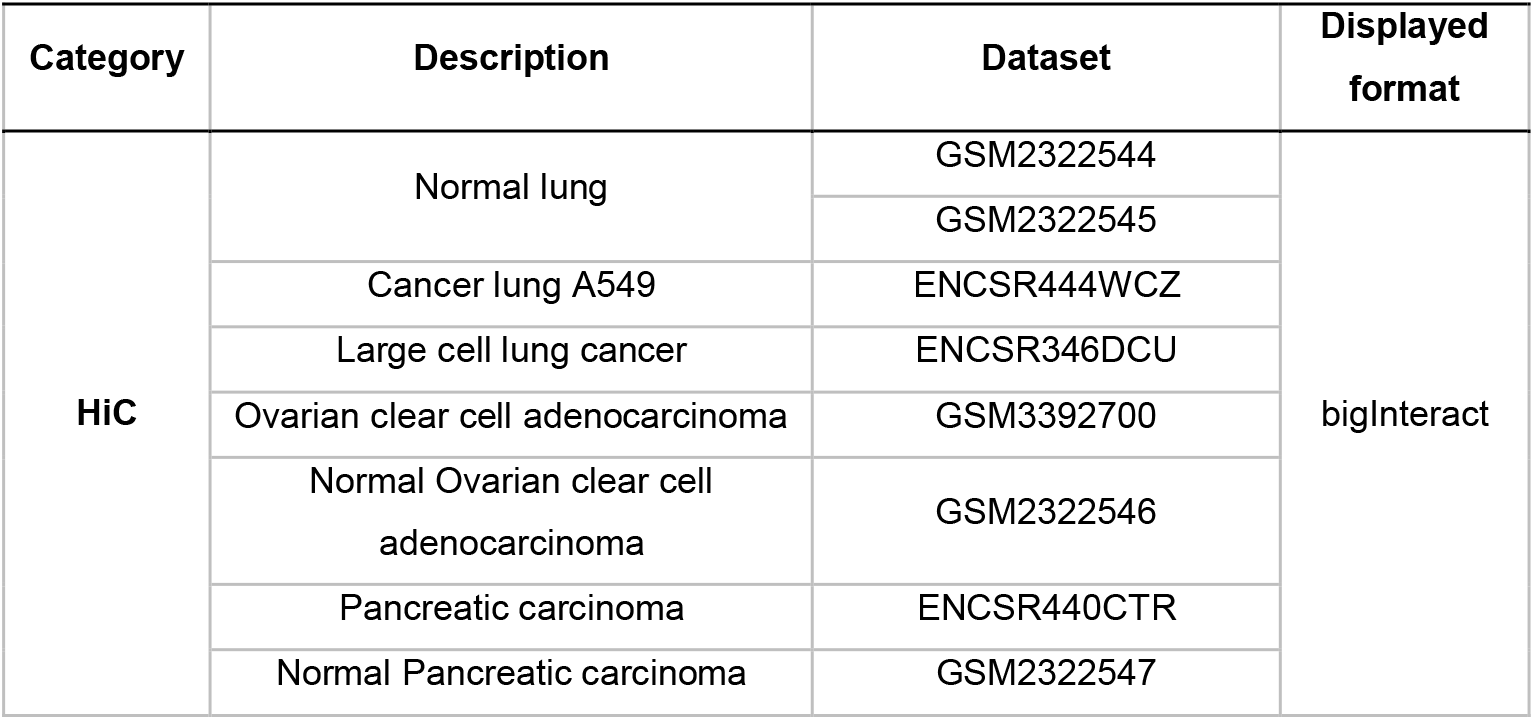
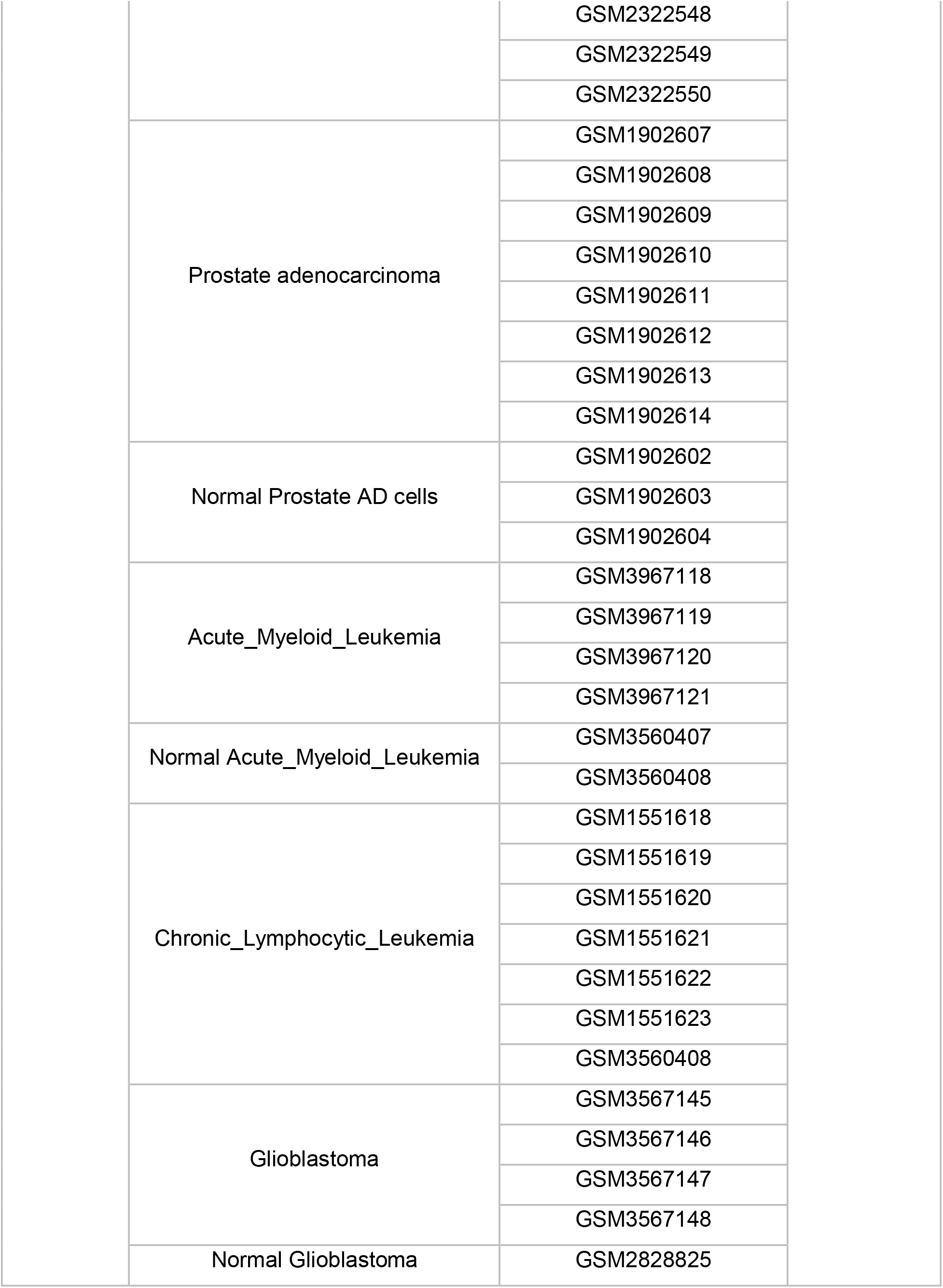
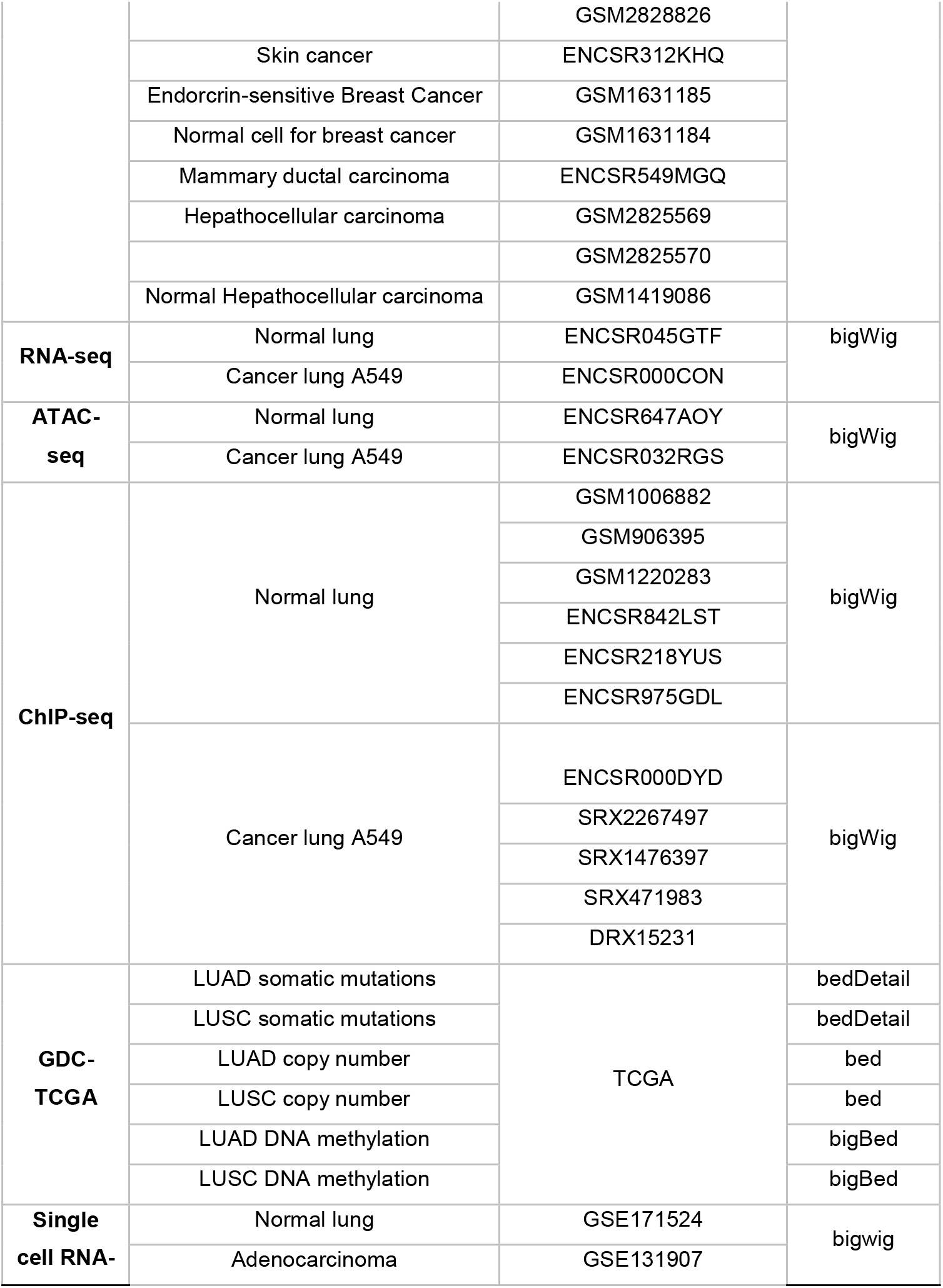
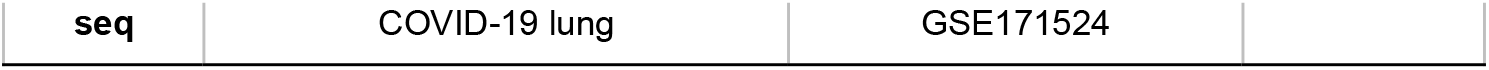
Description of the human lung cancer related multi-omics datasets included in LungCancer3D website

| Category | Description | Dataset | Displayed format |
| --- | --- | --- | --- |
| HiC | Normal lung | GSM2322544 | bigInteract |
|  |  | GSM2322545 |  |
|  | Cancer lung A549 | ENCSR444WCZ |  |
|  | Large cell lung cancer | ENCSR346DCU |  |
|  | Ovarian clear cell adenocarcinoma | GSM3392700 |  |
|  | Normal Ovarian clear cell adenocarcinoma | GSM2322546 |  |
|  | Pancreatic carcinoma | ENCSR440CTR |  |
|  | Normal Pancreatic carcinoma | GSM2322547 |  |

|  |  |  |
| --- | --- | --- |
|  |  | GSM2322548 |
|  |  | GSM2322549 |
|  |  | GSM2322550 |
|  | Prostate adenocarcinoma | GSM1902607 |
|  |  | GSM1902608 |
|  |  | GSM1902609 |
|  |  | GSM1902610 |
|  |  | GSM1902611 |
|  |  | GSM1902612 |
|  |  | GSM1902613 |
|  |  | GSM1902614 |
|  | Normal Prostate AD cells | GSM1902602 |
|  |  | GSM1902603 |
|  |  | GSM1902604 |
|  | Acute_Myeloid_Leukemia | GSM3967118 |
|  |  | GSM3967119 |
|  |  | GSM3967120 |
|  |  | GSM3967121 |
|  | Normal Acute_Myeloid_Leukemia | GSM3560407 |
|  |  | GSM3560408 |
|  | Chronic_Lymphocytic_Leukemia | GSM1551618 |
|  |  | GSM1551619 |
|  |  | GSM1551620 |
|  |  | GSM1551621 |
|  |  | GSM1551622 |
|  |  | GSM1551623 |
|  |  | GSM3560408 |
|  | Glioblastoma | GSM3567145 |
|  |  | GSM3567146 |
|  |  | GSM3567147 |
|  |  | GSM3567148 |
|  | Normal Glioblastoma | GSM2828825 |
|  |  | GSM2828826 |  |
|  | Skin cancer | ENCSR312KHQ |  |
|  | Endocrin-sensitive Breast Cancer | GSM1631185 |  |
|  | Normal cell for breast cancer | GSM1631184 |  |
|  | Mammary ductal carcinoma | ENCSR549MGQ |  |
|  | Hepatocellular carcinoma | GSM2825569 |  |
|  |  | GSM2825570 |  |
|  | Normal Hepatocellular carcinoma | GSM1419086 |  |
| **RNA-seq** | Normal lung | ENCSR045GTF | bigWig |
|  | Cancer lung A549 | ENCSR000CON |  |
| **ATAC-seq** | Normal lung | ENCSR647AOY | bigWig |
|  | Cancer lung A549 | ENCSR032RGS |  |
| **ChIP-seq** | Normal lung | GSM1006882 | bigWig |
|  |  | GSM906395 |  |
|  |  | GSM1220283 |  |
|  |  | ENCSR842LST |  |
|  |  | ENCSR218YUS |  |
|  |  | ENCSR975GDL |  |
|  | Cancer lung A549 | ENCSR000DYD | bigWig |
|  |  | SRX2267497 |  |
|  |  | SRX1476397 |  |
|  |  | SRX471983 |  |
|  |  | DRX15231 |  |
| **GDC-TCGA** | LUAD somatic mutations | TCGA | bedDetail |
|  | LUSC somatic mutations |  | bedDetail |
|  | LUAD copy number |  | bed |
|  | LUSC copy number |  | bed |
|  | LUAD DNA methylation |  | bigBed |
|  | LUSC DNA methylation |  | bigBed |
| **Single cell RNA-** | Normal lung | GSE171524 | bigwig |
|  | Adenocarcinoma | GSE131907 |  |
| seq | COVID-19 lung | GSE171524 |  |

### Data retrieval and processing

#### HiC

Twelve processed HiC datasets involving 9 cancers were obtained from the website of 3D-Genome Interaction Viewer & database (3DIV, http://kobic.kr/3div/download). Each dataset included the signal enrichment over background of each identified HiC loop. For each pairwise comparison between normal tissue cells and tumor cells, we applied fold change threshold of 1.5 to identify the differential loops that specifically exist in each sample.

#### RNA-seq

Raw sequencing fastq files of RNA-seq datasets were downloaded from the ENCODE database, which contain transcriptome data from human lung tissue and lung cancer cells A549. Self-defined Perl pipeline was applied to remove adaptors and low-quality reads with QC < 20 and length < 75bp. Tophat was used to map the reads to hg19 reference genome. The duplicated mapped reads were removed by Picard (version 2.25.0).

These uniquely mapped reads were further processed to identify the differentially expressed genes (DEGs) between human lung and lung cancer cells A549 using Cufflinks (version 2.2.1) with three filters: at least one group FPKM > 1; q < 0.05; log2 fold change > 1.5. These identified DEGs were split into 10 files according to the gradient of q value and fold change and converted to bigWig files. The genes with negative fold change were shown in different shades of blue. Those with positive fold change were shown in different shades of red. The genes that were not differentially changed were shown in grey. For differentially changed genes, the smaller the q values are, the darker the colors are.

#### ATAC-seq

Five paired-end ATAC-seq datasets (3 for A549 cancer cell line, 2 for normal lung tissue) were downloaded from the ENCODE database. TrimGalore (version 0.6.6) was used to trim the reads. Bowtie (version 1.2.2) was used to map the reads and suppress all alignments if more than 1 exist. Picard (version 2.25.0) and samtools (version 1.7) were used to remove the duplicated and unmapped reads with default settings, respectively. The mapped reads were extended to 100bp. Normalized ATAC-seq data was divided into normal lung and cancer cells A549 groups. Data in each group was averaged with Wiggletools (v1.2.2) “mean” function. Differential peaks between normal lung and A549 were identified using bigWigCompare, a tool in Deeptools (v3.3.2), with 200bp bin size. The output bedGraph file were filtered with log2 fold change >1 or <-1 cutoff and were split into two files according to negative or positive fold change. The positive differential peaks were displayed in red and the negative ones were displayed in blue.

#### ChIP-seq

Eleven single-end ChIP-seq datasets (5 for cancer cells A549, 6 for normal lung tissue) were retrieved from the ENCODE and the GEO databases generated from studies of CTCF-binding, H3K27Ac modification, and H3K27me3 modification. Self-defined Perl pipeline was used to filter reads with QC < 20 and length < 50bp. Bowtie, Picard and samtools were used to map and deduplicate reads. The mapped reads were extended to 300bp. The differential peaks were identified and displayed as described above.

#### ScRNA-seq

Three scRNA-seq datasets were downloaded from the GEO database. A scRNA-seq data from normal lung atlas was uploaded within two tracks: ‘10x’, sequencing by 10x Genomics and ‘facs’ sequencing by smartseq2. A scRNA-seq data from lethal COVID-19 atlas was displayed with two tracks: ‘normal’ and ‘covid’. A scRNA-seq data from lung adenocarcinoma was showed with seven tracks: ‘normal lung’, ‘tumor lung’, ‘normal lymph node’, ‘metastatic lymph node’, metastatic brain tissue’, ‘pleural fluid’, and ‘advanced stage tumor lung’. Raw counts data were first separated into different tracks. For the data in each track, we separated cells by their defined cell subtypes, and calculated the FPKM for each gene in each sub-cell-type. The cutoff of FPKM was set to 1. Different cell types in a track were displayed by different colors and the FPKM was displayed by the height of that transcript. All of the above steps were achieved using R (version:4.0) scripts. Then, we obtained chromosome, start and end positions of each transcript from ensembl database using reference ‘gencode.v35lift37.annotation.gtf’ with python3 scripts. For COVID-19 data, we identified DEGs between normal and COVID-19 samples, and for lung adenocarcinoma data, we identified DEGs between normal lung and cancer lung; and normal lymph node with metastatic lymph node. DEG was calculated using log2 fold change of patients sample vs. normal samples with cutoff of 1.5.

#### Somatic mutation (SNP)

The somatic point mutation data was from GDC-TCGA Lung Adenocarcinoma (LUAD) and Lung Squamous Cell Carcinoma (LUSC) cohorts (https://xenabrowser.net/datapages/). Data was preprocessed according to MuSE somatic variant calling workflow. The original alignments were performed using the human reference genome hg38. The tool LiftOver from UCSC Genome browser was used to convert genome coordinates from hg38 assembly to hg19. Mutations with genome coordinates that failed in conversion were excluded from final visualization.

#### Copy number variation

Copy number variation data for LUAD and LUSC were downloaded from GDC-TCGA database (https://xenabrowser.net/datapages/). The original alignments were performed using the human reference genome hg38. The genome coordinates were converted to hg19 assembly using the same method as described above. Currently copy number variation fragments from each sample (patient) are separately presented.

#### Methylation

DNA methylation data for LUAD and LUSC were downloaded from GDC-TCGA database (https://xenabrowser.net/datapages/). Beta value (the ratio of the intensity of the methylated bead type to the combined locus intensity) between 0 to 1 was used to indicate methylation level in these datasets. The beta values from 400 patients were averaged and multiplied by 1000 to fit the bigBed format. Locations of probes were mapped back to hg19 genome using reference files provided by TCGA (https://tcga-xena-hub.s3.us-east-1.amazonaws.com/download/probeMap%2FilluminaMethyl450_hg19_GPL16304_TCGAlegacy).

#### Data display

The mapped reads from RNA-seq, ATAC-seq, and ChIP-seq datasets were converted from bam format to bigwig by genomeCoverageBed and bedGraphToBigWig. To make datasets from different samples with different sequencing depth comparable, normalized bigWig files for RNA-seq, ATAC-seq, and ChIP-seq datasets were generated by bamCoverage (version 3.5.1) using RPKM. Track information (track name, label name, data type, etc.) for the genome browser were stored in file trackDb.ra which was saved in the MySQL root data directory. Track information were uploaded to mysql table hg19.trackDb_NEW with hgTrackDb, a tool provided by UCSC Genome Browser. Tracks were displayed in different file formats, bigInteract, bigWig, bigBed and bedDetail, depending on the data type.

## RESULTS

To demonstrate the applications of LungCancer3D, we utilized the C-MYC region as an example to show the integrated HiC data with other multi-omics data.

### HiC analysis of human normal lungs, lung cancer cells, and other types of cancer

To visualize the dysregulated chromatin loops in human lung cancer, we analyzed the HiC results of normal human lungs and lung cancer cells (A549 cells), respectively. Differential loops between A549 cells and lung cells are identified and displayed (Figure 2A). In addition, we found that there are cancer-specific loops around *C-MYC*, a representative gene (Figure 2B). *C-MYC*, a well-known oncogenic transcription factor, was frequently induced in many types of tumors [28] [29]. Thus, these specifically enriched *C-MYC*-associated loops in lung cancer cells suggest a potential effect of loops on the *C-MYC* expression.

**Figure 2.**
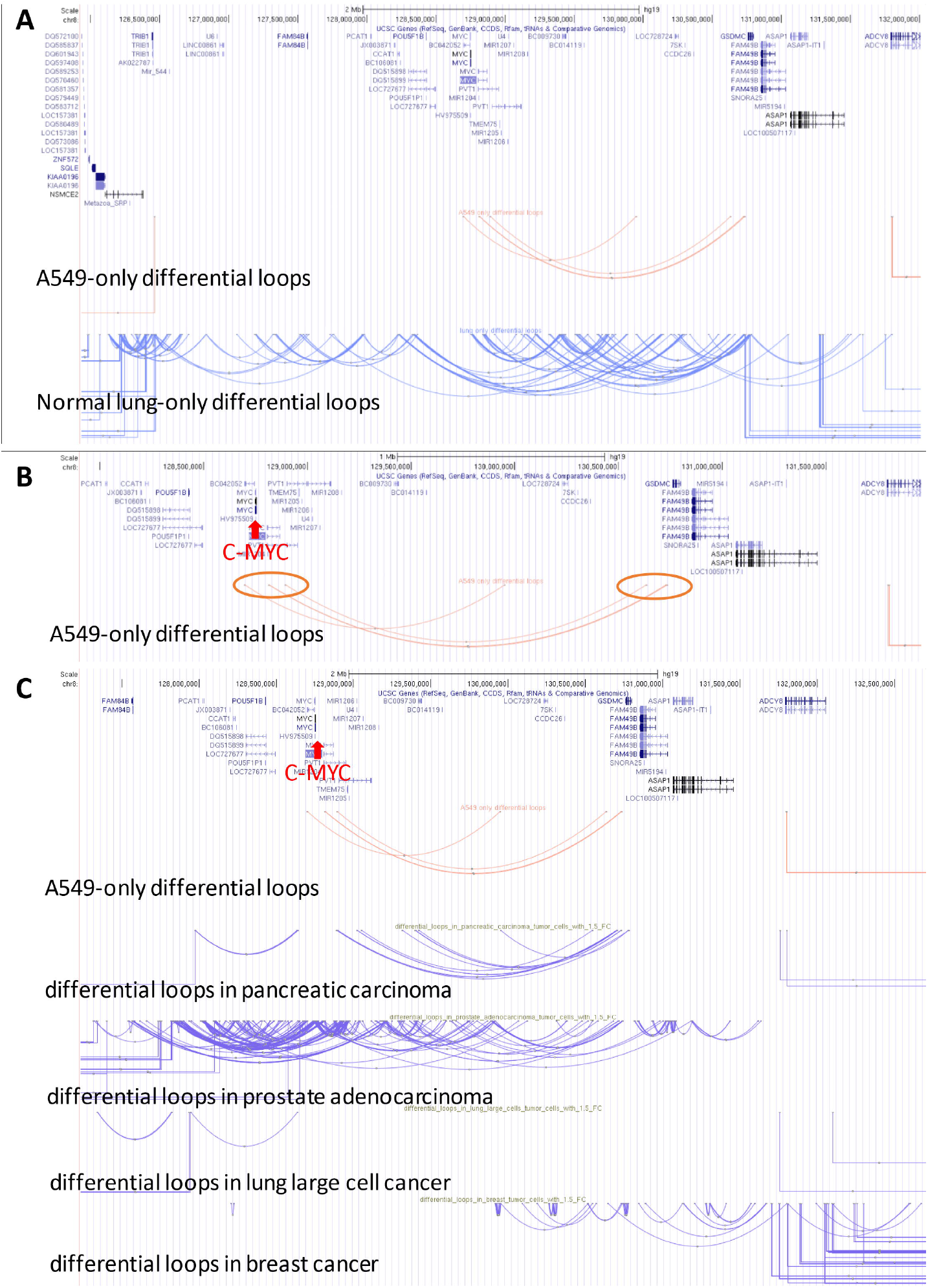
The *C-MYC*-associated loops between normal and tumoral cells. Hi33C results for normal lung cells and A549 cell. (A) Loops that are specifically formed in A549 cells and human lungs. (B) Cancer-specific loops around *C-MYC*. (C) *C-MYC*-associated loops are specifically formed in different tumors.

To investigate if these specifically enriched *C-MYC*-associated loops in human lung cancer are conserved in other types of cancer, we analyzed HiC results in other types of cancer, including lung large cell cancer, breast cancer, pancreatic carcinoma, and prostate adenocarcinoma, and so on. Of note, the *C-MYC*-associated loops showed the tissue-specific patterns though there was a certain similarity among these loops of human lung cancer cells (A549) and pancreatic carcinoma (Figures 2C). Thus, these results indicated that the regulatory mechanism of loops in the *C-MYC* expression was tissue-specific.

### HiC and CTCF ChIP-seq analyses of human normal lungs and lung cancer cells

To examine if there are some chromatin structure proteins (*e*.*g*., CTCF) bound on these loop terminals, we next analyzed the CTCF ChIP-Seq of human normal lungs and lung cancer cells (A549 cells) (Figure 3A). Interestingly, CTCF ChIP-Seq showed that CTCF was bound on the C-MYC-associated loop terminals (Figure 3B). Considering CTCF is a typical chromatin structure protein to form chromatin loops, these analysis results provided the theoretical formation of chromatin loops.

**Figure 3.**
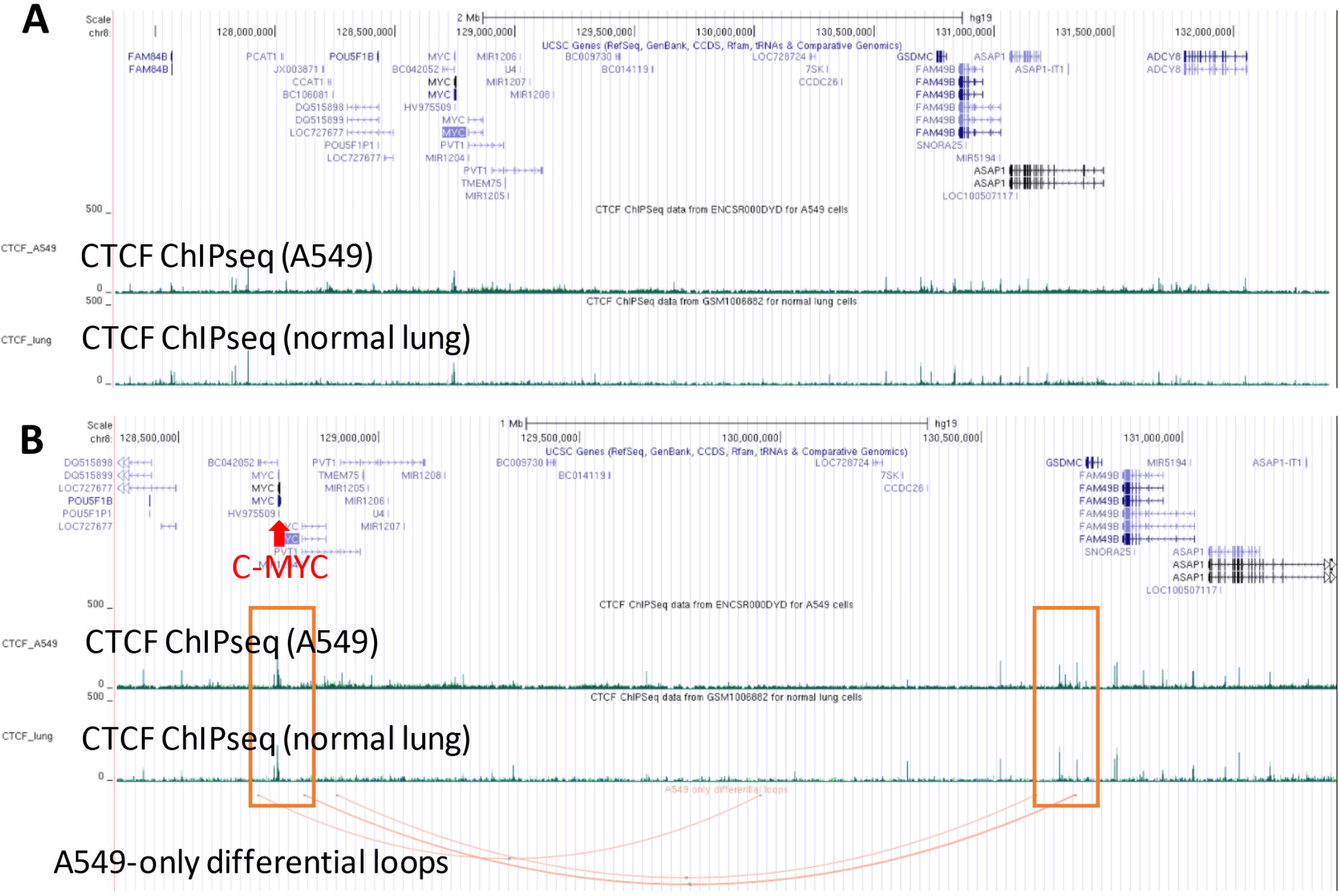
CTCF protein was bound to the terminals of the *C-MYC*-associated loops. (A) CTCF_lung and CTCF_A549 represent CTCF ChIP-seq results for normal lung samples and A549 cells, respectively. The height of each bar indicates normalized coverage of extended reads at each locus. Different colors within each track indicate replicates. (B) CTCF peaks at both ends of loops around C-MYC.

### HiC, CTCF ChIP-seq, and RNA-seq analyses of human normal lungs and lung cancer cells

To investigate if these dysregulated chromatin loops in human lung cancer were associated with the changed gene expression, we further analyze the transcriptome profiles of human normal lungs and lung cancer cells (A549 cells) (Figure 4A). As we expected, the *C-MYC* expression was highly induced in human lung cancer cells, positively associated with the loop (Figure 4B). These results suggest that the potential positive effect of these loops on th*e C-MYC* expression.

**Figure 4.**
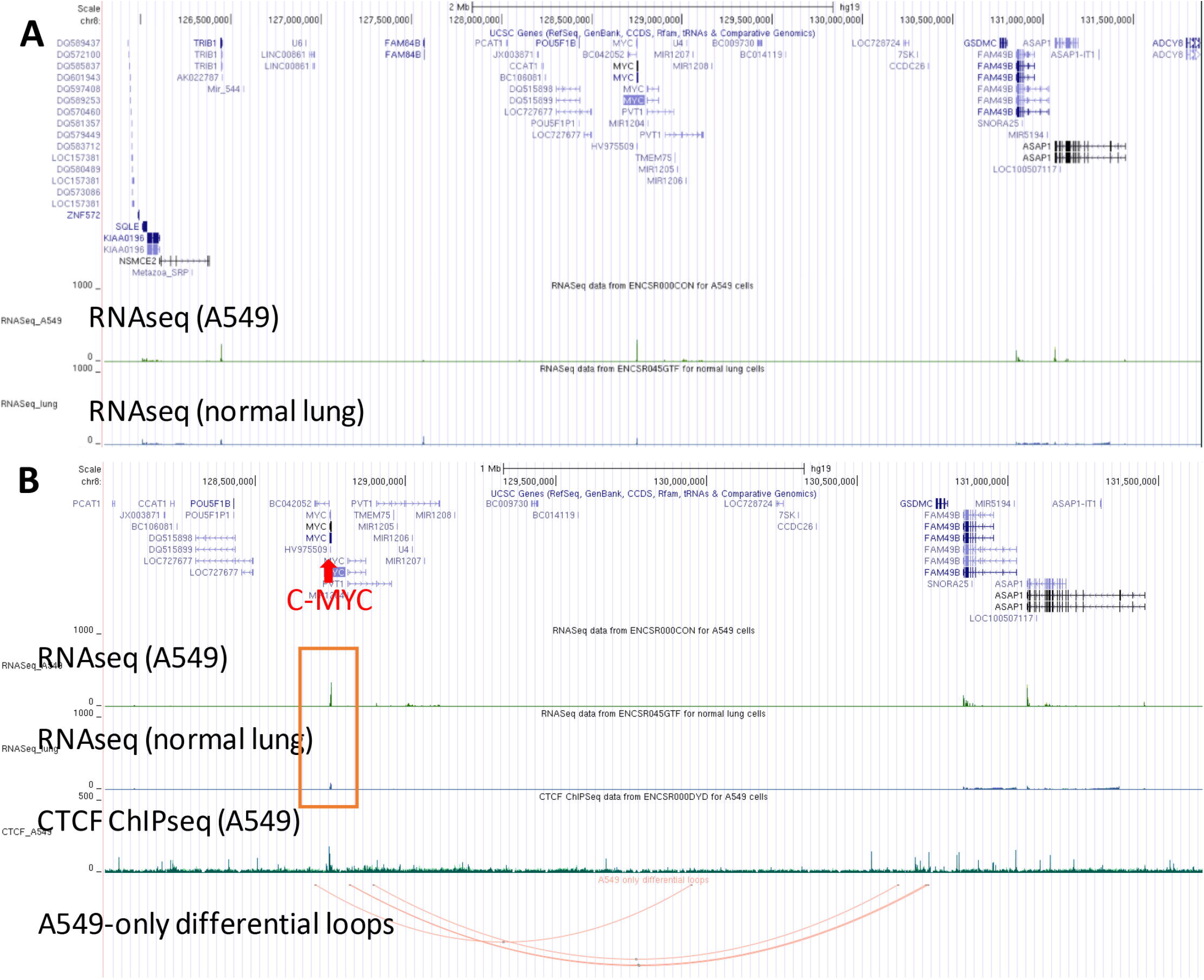
The *C-MYC*-associated loops enriched in human lung cancer cells are positively related to its increased gene expression. (A) RNASeq_lung and RNASeq_A549 track represent RNA-seq results for normal lung samples and A549 cells, respectively. The height of each bar indicates normalized reads coverage at each locus. Different colors within each track indicate replicates. DEG_rnaSeq shows differentially expressed genes, with red for increased expression in lung cancer, grey for decreased expression in lung cancer, height for fold change. (B) C-MYC has significantly increased expression levels in lung cancer with differential loops around it.

### HiC, CTCF ChIP-seq, RNA-seq, and ATAC-seq analyses of human normal lungs and lung cancer cells

To study if open chromatin regions were correlated to loop formations and gene expression, we integrated the ATAC-seq of human normal lungs and lung cancer cells (A549 cells) (Figure 5A). As open chromatin allows functional protein binding and is associated with gene expression, it is expected the chromatin accessibility to be observed where loops form around these changed genes. As expected, we observed that the chromatin accessibility of A549 cells was enhanced compared with normal lung cells at both ends of the *C-MYC*-associated loops (Figure 5B). More interestingly, while A549 cells had ATAC-seq peaks at both ends of the *C-MYC*-associated loop terminals, normal lung cells only had the peaks at one terminal near to the C*-MYC* promoter region, not in the distant region. These results raised the hypothesis that the open chromatin regions, specifically in human lung cancer cells, allowed the structure protein (*e*.*g*., CTCF) to bind and form the new chromatin loop, thereby increasing the *C-MYC* expression.

**Figure 5.**
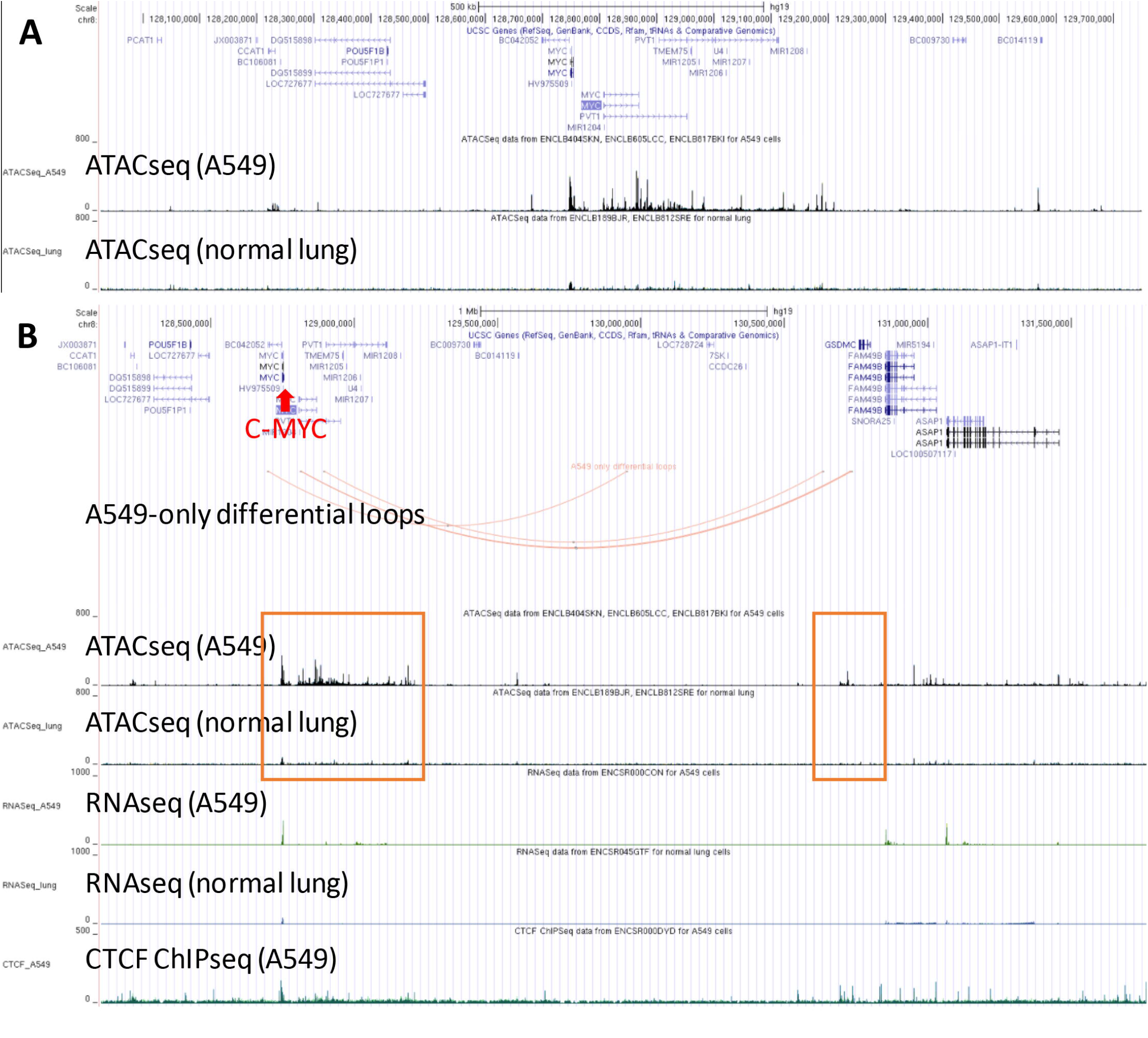
The *C-MYC*-associated loops enriched in human lung cancer cells are in the *de novo* non-promoter open chromatin regions. (A) ATACSeq_lung and ATACSeq_A549 track represent ATAC-seq results for normal lung samples and A549 cells, respectively. The height of each bar indicates normalized coverage of extended reads at each locus. Different colors within each track indicate replicates. (B) At the left end of the differential loop around *C-MYC*, tumor samples have significantly higher chromatin accessibility than normal samples. At the right end, only the tumor sample has some peaks, indicating open chromatin regions.

### HiC, CTCF ChIP-seq, RNA-seq, ATAC-seq, and H3K27ac ChIP-seq analyses of human normal lungs and lung cancer cells

To understand how genes differentially expressed in lung cancer were affected by the specifically enriched loops, we further analyzed H3K27Ac ChIP-Seq of human normal lung cells and lung cancer cells (A549 cell line) since H3K27Ac is a typical marker of transcriptional activation (Figure 6A). At the terminal of *C-MYC*-associated loops near its promoter region, both normal and tumor cells have similar binding profiles of H3K27Ac. However, the H3K27Ac level at that other end of the loop is significantly higher in A549 cells than normal lung cells, suggesting that the loop may bridge the distant enhancer to the *C-MYC* promoter to promote *C-MYC* gene expression (Figure 6B). Furthermore, these H3K27Ac peaks were in these open regions indicated by the ATAC-Seq peaks. These results indicated that the induced *C-MYC* gene expression is due to the enriched loops specifically in lung cancer, which can bridge the distant transcription activation complex to bind on the target gene promoter, resulting in the downstream gene activation.

**Figure 6.**
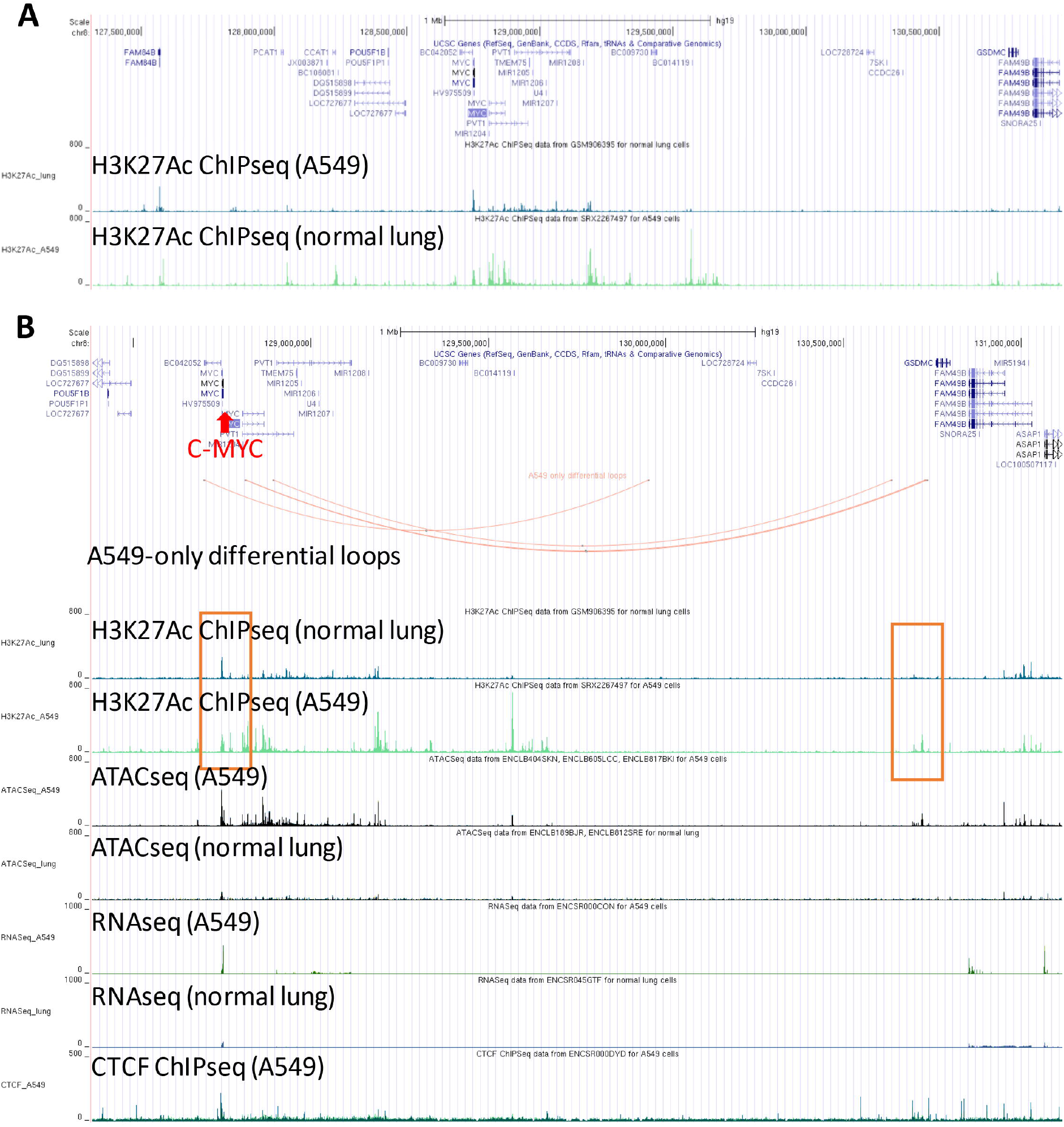
The *C-MYC*-associated loops enriched in human lung cancer cells have the *de novo* binding of H3K27Ac on the non-promoter loop terminal. (A) H3K27Ac_lung and H3K27Ac_A549 represent H3K27Ac ChIP-seq results for normal lung samples and A549 cells, respectively. The height of each bar indicates normalized coverage of extended reads at each locus. (B) At the left end of the cancer-specific loop around C-MYC, normal and tumoral samples have similar H3K27Ac levels. At the right end, only the tumor sample has a peak.

### HiC, CTCF ChIP-seq, RNA-seq, ATAC-seq, H3K27ac ChIP-seq, SNP and methylation analyses of human normal lungs and lung cancer cells

DNA methylation had been considered to be a regulator of loop formation [30]. In detail, loop formation can be decreased with a high methylation level due to the packed chromosome. Therefore, we further analyzed and visualized DNA methylation to investigate the relationship between loop formation and methylation level (Figure 7A). Unexpectedly, the methylation level did not change in lung cancer samples compared to normal lung cells at both ends of *C-MYC*-associated loops (Figure 7B). These results further support the hypothesis that the open chromatin regions, specifically in human lung cancer cells, allowed the structure protein (*e*.*g*., CTCF) to bind and form the new chromatin loop, thereby increasing the *C-MYC* expression.

**Figure 7.**
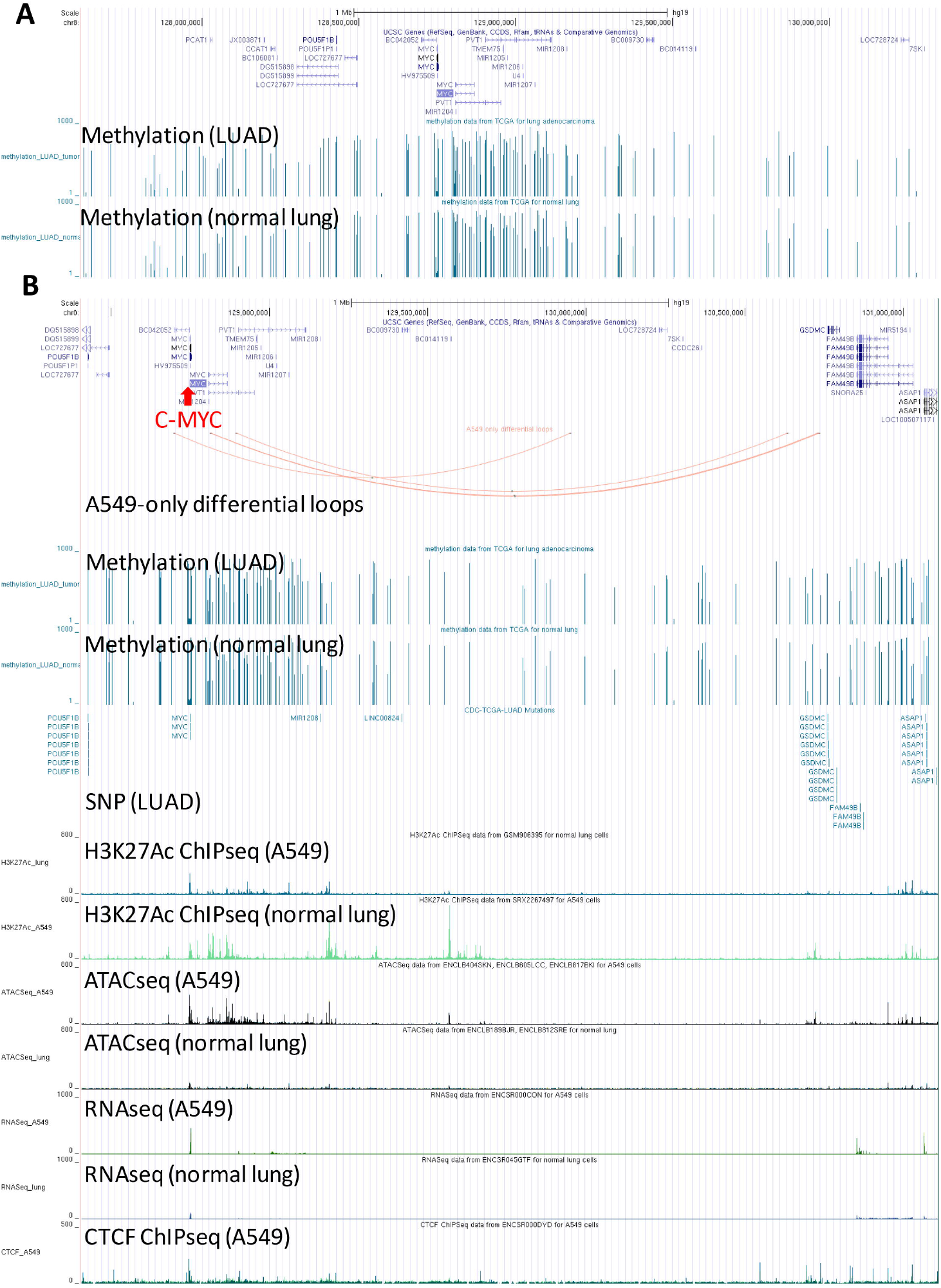
There are no changes in the methylation and SNP in the genomic region of the *C-MYC*-associated loops enriched in human lung cancer cells. (A) The methylation level of tumor cells in LUAD and corresponding normal tissues. The bar height indicates averaged beta value * 1000. (B) Examination of methylation or SNP around the *C-MYC*-related differential loops.

Some long-range interaction between SNPs and targeted genes had been revealed [31]. In addition, some SNPs within the non-coding region may also regulate loop formation, modulating downstream gene expression [32]. Therefore, we also integrated lung cancer-associated SNPs obtained from GDC-TCGA into the browser for users to generate further interpretation (Figure 7B). SNPs on the outer sides of the loop ends for C-MYC located on the coding region for POU5F1B and GSDMC, indicating an interplay between these SNPs and loops.

### HiC, CTCF ChIP-seq, RNA-seq, ATAC-seq, H3K27ac ChIP-seq, and single-cell RNA-seq analyses of human normal lungs and lung cancer cells

As loops may regulate gene expressions as shown in the bulk RNA-seq result, we wonder if they are associated with gene expressions in certain cell types. We also wonder whether COVID-19 affection will induce lung cancer-related gene expression. Therefore, we analyzed single-cell data for human normal lungs, lung adenocarcinoma, lungs with COVID-19 infection (Figure 8A). By comparing gene expression between lung cancer and normal lungs, we found that genes related to lung cancer, such as *C-MYC*, had much higher expressions in many types of cells (including EPC cells, smooth muscle cells, Pericytes cells) of lung cancer patients (Figures 8B). However, the *C-MYC* gene did not have a higher expression in COVID-19 infected patients (Figure 8C), which indicates that the potential impact of COVID-19 infection on lung cancer may not be correlated to C-MYC.

**Figure 8.**
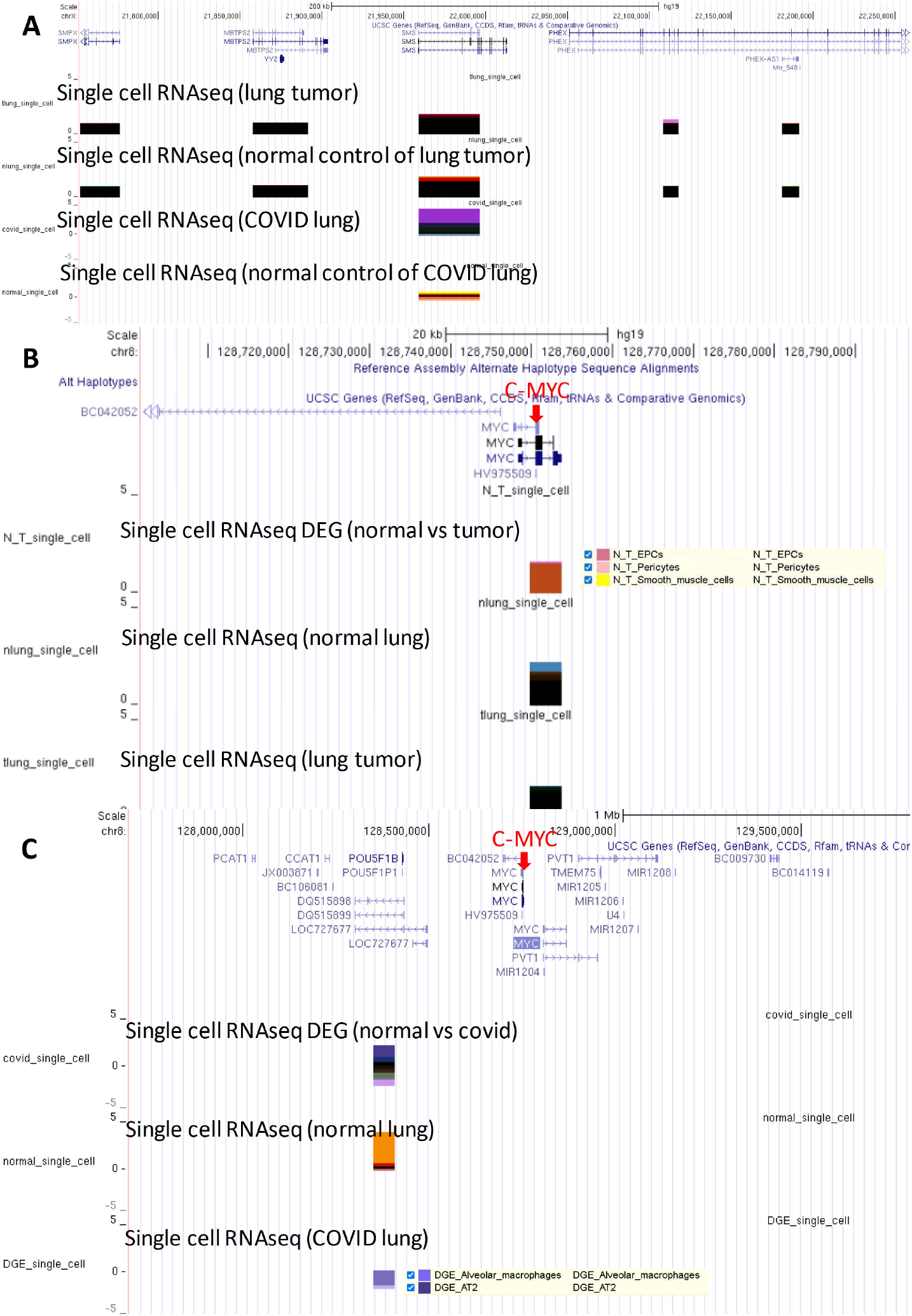
single-cell transcriptome is correlated with cancer-specific loops. (A) from top to the bottom, DEGs of normal lung, lung cancer and COVID-19 patients single-cell samples. Height indicates fold change and color represents cell type. (B) C-MYC is a differentially expressed gene in many types of cells of lung cancer patients. (C) C-MYC is not a DEG in any types of cells of COVID-19 patients.

## DISCUSSION

Here we built a UCSC browser-based website database, LungCancer3D, to meet the increasing needs for 3D chromatin architecture studies and multi-omics data analyses. The website database currently displays 10 different multi-omics data, including HiC-loops, RNA-seq, snRNA-seq, ATAC-seq, CTCF ChIP-seq, H3K27AC ChIP-seq, H3K27me3 ChIP-seq, DNA methylation, CNVs, and SNP mutations. We utilized the *C-MYC* gene in lung cancer cells as an example to demonstrate the directly visualizing these multiple omics data and found that its induced expression in lung cancer cells might be caused by the *de novo* loop formation bridging the distant enhancer to bind on its promoter region.

The most significant advantage of multi-omics analyses is to explicate that the differential changes (*e*.*g*., H3K27Ac ChIP-seq peak) identified in one omics data with the systematic considering of other layers of molecular changes, helping reveal the mechanisms unbiasedly. For example, the increased H3K27Ac binding (H3K27Ac ChIP-seq peak) at non-coding genome regions is frequently neglected by researchers since they are hard to link to gene expression regulation. However, with the HiC-identified chromatin loops in these regions and the nearby ATAC-Seq and CTCF ChIP-seq peaks, the increased H3K27Ac binding at non-coding genome regions can be linked to the target gene, which is C-MYC. Therefore, the primary function of our website of the current version is to help users directly visualize multi-omics data.

The *C-MYC* gene is a typical oncogene in many types of cancer [33]. A previous study illuminated that the duplication of the C-MYC enhancer region caused the upregulation of C-MYC [29]. Unlike these previous findings, our observation from LungCancer3D showed that an enhancer-promoter interaction might be due to the formation of a new loop that only exists in lung cancer, which brought an enhancer region to bind the *C-MYC* promoter. We also observed that the *de novo* non-promoter loop terminal in lung cancer with increased H3K27Ac histone bindings had relatively decreased bindings of H3K27me3 histone. Thus, histone methylation and acetylation may correlate with loop formation. We also found some lung cancer somatic mutations near *C-MYC*. These SNPs are in the genomic region of the *POU5F1B* and *GSDMC* genes. Previous studies report that Caspase-8 cut GSDMC at D365 amino acid in cancer [34] [35], thus remodeling the chromatin architecture. Since the loops formed in lung cancer are different from those in other cancers, such as breast cancer or duct cancer. It suggested that a site that represses the formation of this lung cancer loop may specifically target lung cancer cells. A closer study on these SNPs may help researchers identify a novel site for target therapy.

In the future, besides paying more attention to the study of 3D chromatin architecture changes at the level of multiple omics, we will continue optimizing our website. We will increase the size of each omics data by combining and analyzing more datasets and allowing users to upload their data to our website in either raw data format or processed data format. In addition, we will add functions that analyze the differential gene expression, differential peaks, and differential loops and allow users to download them and other data. The goal is to construct a website database with tools for users to generate differential data in multiple omics and a comprehensive dataset for lung cancer study.

## DATA AVAILABILITY

LungCancer3D is publicly available at http://www.lungcancer3d.net to all users. All these datasets were downloaded from online databases and were described in Table 1.

## CONFLICT OF INTEREST

The authors declare that they have no conflict of interest.

## FUNDING

This work was supported by grants to JL from the National Natural Science Foundation of China (82172899), “Oncology Therapy” Project of Hangzhou Medical Peak Discipline, Interdisciplinary Research of Young Scholars of Zhejiang University International Campus (Li Ye Foundation), and startup funding of Zhejiang University.

## ACKNOWLEDGEMENTS

We want to thank the UCSC Genome Bioinformatics Group for downloading, modifying, and rehosting the browser. In addition, we are grateful to all researchers who have generated and shared their multi-omics data. We also thank the help of other group members in Dr. Jian Liu’s lab.

## AUTHORS’ CONTRIBUTIONS

JL conceived, designed, and supervised the research; XW, AJ, JW contributed to the conception and design of the study; XW, AJ, JW, SS built up the website; AJ, JW, XW, SS collected and analyzed metagenome data and uploaded them to the website; SS, AJ, JW, XW, JL prepared the manuscript, JL revised the manuscript; SM, XC, BX, SZ, YX, QT tested the website for optimization.

## Notes

### Competing Interest Statement

The authors have declared no competing interest.

